# Testosterone differentially modulates the display of agonistic behavior and dominance over opponents before and after adolescence in male Syrian hamsters

**DOI:** 10.1101/2025.03.31.646499

**Authors:** Arthur J. Castaneda, Conner J. Whitten, Tami A. Menard, Cheryl L. Sisk, Matthew A. Cooper, Kalynn M. Schulz

## Abstract

The current study investigated the influence of testosterone on agonistic behavior and dominance over an opponent before and after adolescence in male Syrian hamsters (*Mesocricetus auratus)*. We hypothesized that testosterone-dependent modulation of agonistic behavior would be greater following adolescent development. To test this hypothesis, prepubertal (14 days of age) and adult subjects (52-62 days of age) were gonadectomized and immediately implanted with testosterone or vehicle pellets. Fourteen days later, agonistic behavior was assessed in a neutral arena with age-matched testosterone-treated opponents. Flank marking was also assessed separately in response to male odors alone. Our hypothesis predicted that testosterone would modulate agonistic behavior and dominance over an opponent in adult but not in prepubertal subjects, however, only flank marking behavior followed the predicted data pattern. During both social interaction and scent tests, testosterone increased flank marking behavior in adults, but failed to increase flank marking in prepubertal subjects. Contrary to our predictions, testosterone treatment increased prepubertal subject attacks, decreased submissive tail-up displays, and facilitated prepubertal subject dominance over opponents. In adults, testosterone increased paws-on investigation and flank marking during social interactions. Taken together, these data indicate that some, but not all aspects of agonistic behavior are sensitive to the activational effects of testosterone prior to adolescence, and that activational effects of testosterone differ substantially between prepubertal and adult males. Our results may have implications for early pubertal timing and increased risk for externalizing symptoms and aggressive behavior in humans.

**Highlights:**

- Testosterone increased attacks and decreased submissive displays in prepubertal males
- Testosterone increased dominance over opponents in prepubertal males
- Prepubertal males displayed more attacks and submissive behaviors than adults overall
- Testosterone increased flank marking behavior only in adult males

## 1.0 Introduction

Dominance hierarchies are formed in virtually every animal species ranging from insects to mammals. Social dominance confers an important advantage for acquiring resources such as food (Johnston, 1970) and mating opportunities (Huck & Lisk, 1985; Wallen, 1982; Wallen & Goy, 1977). A dominance hierarchy is achieved when one animal threatens, chases, or attacks while the opponent displays defensive or submissive behavior (Chase et al., 2002). Syrian hamsters are an excellent animal model for the study of social communication and dominance relationships. Upon initial contact, adult male hamsters progress through a series of behaviors that include initial approach, offensive and defensive postures, attacks in attempt to bite, and flank marking. Flank marking occurs when hamsters rub specialized dorsolateral flank glands onto surfaces in their environment (Drickamer et al., 1973; Ferris, 1996; Ferris et al., 1987; Whitsett, 1975). When hamsters are tested in pairs, individual that exhibits more aggressive displays (e.g., attacks and offensive postures), and fewer submissive displays (e.g., defensive and tail-up postures), is typically considered to be dominant. These dominance relationships are formed quickly and remain relatively stable across time (K. C. De Lorme & Sisk, 2013; Ferris et al., 1987; Grieb et al., 2021; Morrison et al., 2012; Whitten et al., 2023). Across repeated pairings, the dominant male’s overt aggression decreases, whereas flank marking increases, suggesting that flank marking may serve to maintain dominance relationships and reduce the need for continued overt aggression (Ferris et al., 1987). However, given that dominance hierarchies form quickly during a social encounter (Ferris et al., 1987; Goldman & Swanson, 1975; Payne & Swanson, 1970), flank marking may also establish an individual’s dominance status by working in concert with other aggressive displays.

Male social interactions change dramatically during adolescence. For example, juvenile male Syrian hamsters attack opponents at higher frequencies than adults (Cervantes et al., 2007; Romeo et al., 2003; Wommack et al., 2003), and tend to target attacks toward the head and cheeks of opponents, whereas adult males target attacks toward the lower belly and flank area (Cervantes et al., 2007; Pellis & Pellis, 1988a, 1988b; Wommack et al., 2003). Pubertal testosterone does not seem to underly the adolescent decrease in attack frequency (Romeo et al., 2003) or adolescent changes in the bodily target of attacks (Smith et al., 1996). However, it is unknown whether pubertal testosterone modulates the full repertoire of agonistic behavioral displays, including those communicating submissive status.

The current study investigated the effects of testosterone on agonistic behavioral displays and dominance over an opponent in juvenile and adult males. Flank marking behavior was also assessed in response to odors contained in adult male bedding. By comparing the effects of gonadectomy and testosterone replacement before and after adolescent development, we sought to determine whether activational effects of testosterone are present at pubertal onset, or whether behavioral neural circuits acquire responsiveness to testosterone during adolescence. Our results indicate that prepubertal and adult males display unique agonistic behavioral profiles in response to testosterone.

## 2.0 Material and Methods

### 2.1 Experimental Design

See Figure 1 for a timeline of experimental procedures. A two-factor between subjects design was employed to assess the effects of Age (prepubertal vs. adult) and Hormone (testosterone vs. vehicle) on agonistic behavior. Prepubertal subjects were castrated at 14 days of age, and adult subjects were castrated between 52-62 days of age. At the time of castration, subjects were implanted with a beeswax pellet containing testosterone propionate (T) or a control (vehicle) beeswax pellet, resulting in 4 experimental groups: Prepub+0, Prepub+T, Adult+0, and Adult+T. Two weeks later, prepubertal (28 days of age) and adult subjects (66-76 days of age) underwent two behavioral tests separated by a 10-minute resting period: 1. A scent test to assess flank marking behavior in response to the soiled bedding of unfamiliar adult males, and 2. A social interaction test to assess aggressive and submissive behavioral displays during a social interaction with an opponent. Presentation of the scent and social interaction tests was counterbalanced. Opponents were age- and weight-matched within 5g to a subject and only tested with one subject male. To account for age-dependent differences in circulating testosterone between prepubertal and adult opponents, all opponents were castrated and T-treated one week prior to social interactions.

**Figure 1.**
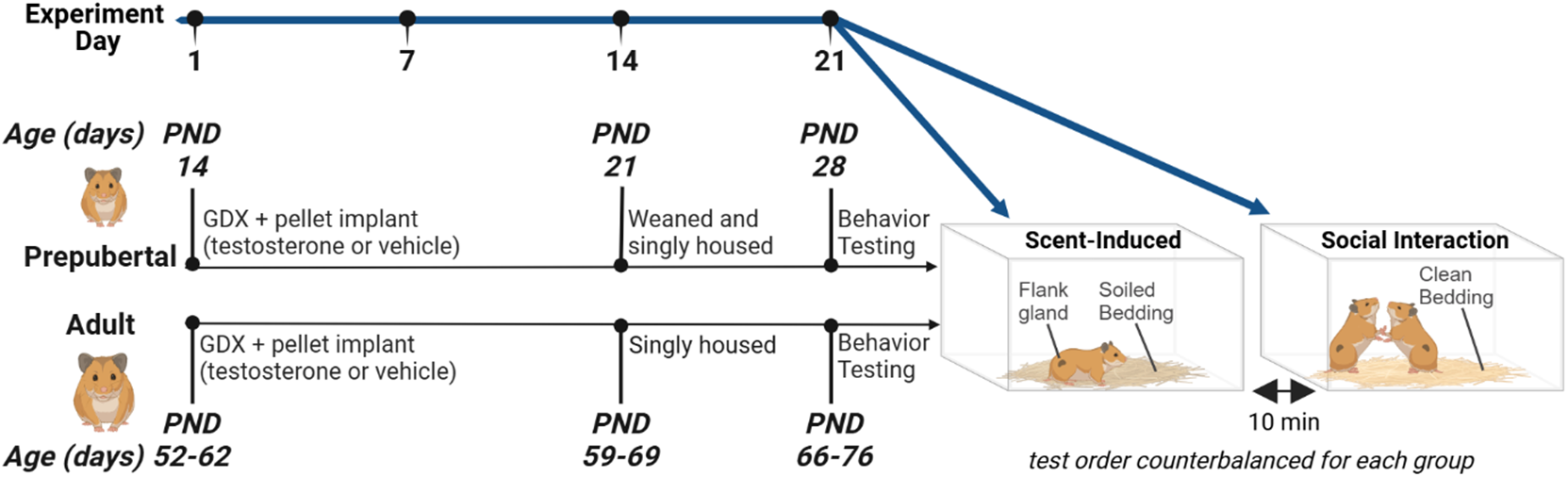
Experimental Design. Prepubertal and adult subjects were gonadectomized (GDX) and immediately implanted with testosterone (T) or vehicle pellets, resulting in 4 groups: Prepub+0, Prepub+T, Adult+0, and Adult+T. Subjects were behavior tested two weeks later, and all subjects were singly housed for 1 week prior to scent-induced flank marking tests and social interaction tests. In the scent-induced flank marking test, subjects were placed into a testing aquarium filled with the soiled bedding of gonad-intact adult males and the number of flank marks were quantified. In the social interaction test, subjects were age- and weight-matched with T-treated male opponents, and the number of aggressive and submissive behavioral displays were quantified.

### 2.2 Animals

All animals were housed in a 14 h light 10 h dark reverse light schedule (lights off at 1200 h EST) to minimize the variation in circadian rhythms and maintain an aggressive reproductive state (Landau, 1975). Food (Teklad Rodent Diet No. 8640, Harlan) and water were available *ad libitum*. Animals were treated in accordance with the NIH Guide for the Care and Use of Laboratory Animals, and all protocols were approved by the Michigan State University Institutional Animal Care & Use Committee.

#### 2.2.1 Prepubertal and Adult Subjects

Twenty-three prepubertal subjects were bred in laboratory from Harlan Sprague-Dawley stock. Adult subjects (n = 23) ranging in age from 50-60 days were received from Harlan Sprague-Dawley (Indianapolis, IN) and group housed on arrival. Prepubertal subjects were weaned at 21 days of age, at which time they were single housed. To avoid potential cohort effects, prepubertal and adult subjects were single housed and behavioral tested in parallel. Specifically, all subjects were single housed (37.5 x 33 x 17 cm) for 7 days prior behavioral testing.

#### 2.2.2 Prepubertal and Adult Opponents

Prepubertal opponent males arrived with their dams and littermates from Harlan Sprague-Dawley at 17, 18, and 19 days of age, and were weaned/single housed at 21 days of age. Adult opponents arrived at 50-60 days of age and were group housed upon arrival. Like subjects, all opponents were single housed for 7 days prior to social interaction tests.

### 2.3 Surgical Procedures

#### 2.3.1 Castration and Testosterone Administration

Castrations and testosterone implants were performed in one surgical procedure under isoflurane anesthesia (oxygen with 3-5% isoflurane, flow rate 0.7 to 1 L/min). Males were administered a subcutaneous injection of the analgesic buprenorphine (0.05 mg/kg) prior to surgery. The testes were pulled through bilateral scrotal incisions, and the testicular veins were tied with suture silk before removal of the testes. The incisions were closed with 9mm autoclips (Becton Dickinson, 427631). Beeswax pellets containing testosterone propionate (T) or vehicle (vehicle) were inserted subcutaneously through a 5 mm incision made on the dorsal midline between the scapulae of the animal, and the incision was closed with an autoclip.

#### 2.3.2 Testosterone Propionate Beeswax Pellets

T dissolved in 95% ethanol was added to melted beeswax (Sigma-Aldrich) at a concentration of 0.05 mg testosterone / 1.0 mg beeswax. A metal punch tool was used to extract pellets from the hardened beeswax mixture. The dose of testosterone administered to animals was determined by the weight of the beeswax pellet, and all pellet weights were verified prior to implantation. Two weeks prior to behavioral testing, subjects were castrated and implanted with a 100 mg beeswax pellet containing 5.0 mg testosterone or beeswax alone (vehicle). In contrast, one week prior to behavioral testing, all opponents were castrated and implanted with a 50 mg pellet containing 2.5 mg T. Testosterone levels were matched between prepubertal and adult opponents to reduce the likelihood that any behavioral differences between prepubertal and adult subjects were due to variations in the circulating testosterone levels of the opponents (Evans & Brain, 1974; Payne, 1974; Solomon et al., 2009). Pilot studies determined that beeswax pellets decrease their release of testosterone over time. Therefore, we adjusted the testosterone dose administered to opponents to account for their shorter time interval between surgical implants and behavioral testing. This adjustment ensured that both subjects and their opponents would have physiological levels of circulating testosterone during social interactions. Plasma testosterone concentrations were confirmed via radioimmunoassay (Table 2).

### 2.4 Behavioral Testing and Scoring

Behavioral testing began one hour into the dark phase of the light cycle. Prepubertal and adult subjects were tested sequentially in two counterbalanced conditions: (1) A 10 min social interaction test in which subjects were paired with an age- and weight-matched T-treated opponent, and (2) a 10 min scent test in which flank marking behavior was observed in response to the soiled bedding of male conspecifics. Soiled bedding was collected from the cages of gonad-intact adult males and stored in an air-tight container. These males were not part of the current study and unfamiliar to all study animals. One cup of soiled bedding was scattered across the floor of the glass aquarium (61 x 32 x 31 cm) immediately before the scent test. For social interaction tests, one cup of clean bedding was scattered across the floor of the aquarium before subjects and opponents were simultaneously placed inside. To ensure that opponents were naïve, they received only one social interaction test. The walls and floor of the aquarium were cleaned thoroughly with 70% ethanol and allowed to dry completely between tests, and all tests were video recorded and scored by one experimenter blind to treatment condition (Table 1).

**Table 1.** Behavioral Scoring Criterion.

| Behavior | Scoring Criterion |
| --- | --- |
| **Flank Mark** | The subject rubs their dorsolateral flank gland against the testing chamber wall. |
| **Close Contact** | The physical distance between subjects and opponents is within 3 cm. |
| **Aggressive Displays:** |  |
| *Attack* | The subject moves quickly toward the opponent and attempts to bite. |
| *Paws-on Investigation* | The subject applies forepaw pressure onto the opponent's back while actively investigating/sniffing. The opponent stands still during paws-on investigation. |
| *Offensive Posture* | The subject stands upright on their hind legs facing the opponent. The opponent must also be displaying an offensive or defensive posture. |
| **Submissive Displays:** |  |
| *Tail-up Posture* | The subject raises their tail (above parallel to the floor) and rounds their back upward (kyphosis). This posture can be assumed when walking or standing still. |
| *Defensive Posture* | The subject twists their upper body toward the opponent with one or both forepaws outstretched to ward off physical contact. |

#### 2.4.3 Social Interaction Test: Subject Aggression & Win Index Score

A win index was calculated to investigate whether subjects dominated their opponent during the social interaction. First, an aggression score was calculated separately for subjects and opponents by subtracting an individual’s average number of submissive behavior displays (tail-up + defensive postures /2) from their average number of aggressive behavior displays (attacks + paws-on + offensive postures/3). The subject’s win index was then computed by subtracting the opponent’s aggression score from the subject’s aggression score. Subjects with an index greater than zero were more dominant than their opponent during the social interaction (winner), and subjects with an index less than zero were more submissive than their opponent (loser).

### 2.5 Blood Collection and Testosterone Radioimmunoassay

Subjects were weighed and administered an overdose of sodium pentobarbital (130 mg/kg intraperitoneal). Blood was collected via cardiac puncture into EDTA-coated tubes and centrifuged at 4 ℃. Plasma was removed and stored at -20 ℃ until radioimmunoassay. Testosterone concentrations were measured in duplicate 50µL samples within a single assay using the Coat-A-Count Total T Kit (Diagnostic Products, Los Angeles, CA). This assay has been previously validated (Parfitt et al., 1999). The intra-assay CV was 9.2%, and the lower limit of detectability was 0.1 ng/ml.

### 2.6 Flank Gland Measurement

Flank gland diameter was measured immediately following blood collection. The flank gland (also called the flank organ) is a slightly raised and oval shaped collection of sebaceous scent glands located bilaterally on the dorsolateral flanks. The flank gland increases in diameter and center pigmentation during puberty and is highly responsive to circulating androgen levels in adulthood (Algard et al., 1966; Hamilton & Montagna, 1950). We investigated whether flank gland responsiveness to testosterone differs between prepubertal and adult subjects. The hair overlying the subject’s right flank gland was shaved before calipers were used to measure the largest diameter of the palpable bulk in millimeters. A central region of dark pigmentation was observed only in T-treated subjects.

### 2.7 Sample Size and Experimental Attrition

Twenty-three prepubertal and 23 adult subjects underwent behavioral testing procedures, although the following issues resulted in data loss and/or exclusion from analysis. Radioimmunoassay confirmed the failure of a testosterone implant in one prepubertal subject, and their data were excluded from all analyses. Technical problems with video recording equipment resulted in the loss of two subjects’ social interaction test data, and one subject’s scent-induced test data. Finally, one subject’s social interaction test data were excluded from analysis because they displayed abnormally elevated levels of flank marking behavior. Specifically, this subject’s display of 61 flank marks was 4.97 standard deviations above the mean for all subjects, and more than two times greater than the next highest value of 28 flank marks. Thus, group sizes for analyses ranged between 9-12 subjects.

### 2.8 Data Analysis

The Kolmogorov-Smirnov test of normality was conducted for each dependent measure. Normally distributed data were analyzed by 2-factor ANOVA or independent t-tests. Parametric test results were interpreted using p values, estimates of effect size, and confidence intervals. For ANOVA, effect size was estimated using partial Eta squared values (small effect, ηp^2^ = 0.01-0.059; moderate effect, ηp^2^ = 0.06-0.139; large effect, ηp^2^ = 0.14 and greater), whereas t-tests were followed by Cohen’s D estimate of effect size (small effect, d = 0.2; moderate effect, d = 0.5; large effect, d= 0.8). Non-normally distributed data were analyzed by the Kruskal-Wallis nonparametric test. Statistical significance was considered *p* `< 0.05.`

## 3.0 Results

### 3.1 Testosterone Radioimmunoassay

Plasma testosterone concentrations in T-treated subjects fell within the physiological range of 2-7 ng/ml found in gonad-intact adult male Syrian hamsters (Sisk and Turek, 1985). Beeswax T-pellets yielded similar levels of plasma testosterone in prepubertal and adult subjects (Table 2; *Mean Difference* = 0.52 ng/ml, CI 95% [-0.72 - 1.76 ng/ml], *t* (1, 20) = .386, p = .39, d = .38). Similarly, no differences in plasma testosterone concentrations were found between the opponents that were paired with subjects (Table 2; *F* (3, 38) = 1.10, p = .36, ηp^2^ = .08). We also assessed potential differences in plasma testosterone between subjects and their opponents. For prepubertal males, no significant differences in testosterone concentrations were found between testosterone-treated subjects and their opponents (*Mean Difference* = 0.80 ng/ml, CI 95% [-0.43 - 2.04 ng/ml], *t* (1, 18) = 1.37, p = .19, *d* = .61). For adult males, a marginally significant difference was found between testosterone-treated subjects and their opponents (subjects > partners; *Mean Difference* = 0.86 ng/ml, CI 95% [-0.04 - 1.75 ng/ml], *t* (1, 22) = 2.03, p = .059, *d* = .83). Prepubertal and adult subjects treated with vehicle pellets displayed testosterone concentrations below the lower limit of assay detectability and were not analyzed.

**Table 2.** Mean plasma testosterone concentrations in prepubertal and adult subjects and their opponents.

| Subject Treatment Group | Subjects | Opponents |
| --- | --- | --- |
|  | Testosterone (ng/ml) | Testosterone (ng/ml) |
| **Prepub+0**<br>(n = 11) | *undetectable* | 3.39 (0.51) |
| **Prepub+T**<br>(n = 10) | 3.98 (0.46) | 3.18 (0.37) |
| **Adult+0**<br>(n = 9) | *undetectable* | 2.73 (0.28) |
| **Adult+T**<br>(n = 12) | † 3.47 (0.37) | 2.61 (0.20) |

† indicates a marginally significant difference ( $p < .1$ ) between T-treated adult subjects and their opponents. No differences in plasma testosterone levels were found between T-treated prepubertal and adult subjects. Likewise, no differences were found between the opponents of any subject treatment group.

### 3.2 Body Weight and Flank Gland Diameter

As expected, body weights were significantly greater in adults than in prepubertal subjects (Figure 2A; *F (*1, 41) = 671.42, p < .001, ηp^2^ = 0.94). testosterone treatment did not significantly impact body weight (*F* (1, 41) = 2.47, p = .124, ηp^2^ = .06), nor did Age and Hormone interact to influence terminal body weights (*F* (1, 41) = .464, p = .500, ηp^2^ = .01).

**Figure 2.**
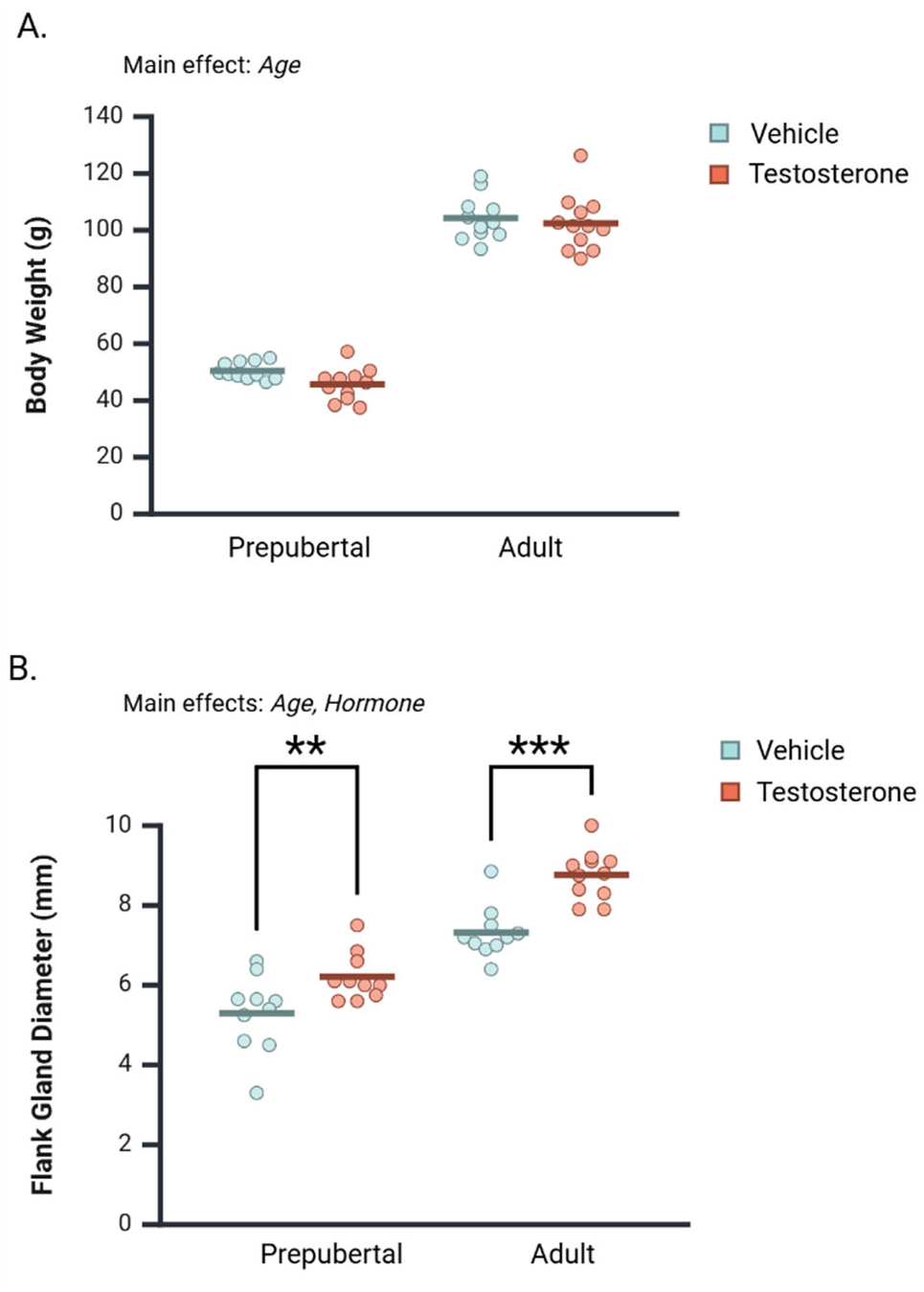
Mean body weight and flank gland diameter in castrated prepubertal and adult subjects that received 2 weeks of testosterone or vehicle treatment. **A.** Testosterone treatment did not significantly affect body weight in juvenile or adult subjects. **B**. Adults exhibited greater flank gland diameters overall, and testosterone administration significantly increased flank gland diameter in both prepubertal and adult subjects. Circles represent individual values, and the bars represent the group mean. **\*\*** denotes p < .01; **\*\*\*** denotes p < 0.01.

Flank gland diameter was measurable in vehicle- and testosterone-treated subjects, although only testosterone-treated subjects exhibited a darkly pigmented central region. Both Age and Hormone impacted flank gland diameter (Figure 2B). Specifically, testosterone-treatment significantly increased flank gland diameter (*F* (1, 37) = 27.33, p < .001, ηp^2^ = .43), and adults exhibited greater flank gland diameters than prepubertal subjects (*F* (1, 37) = 102.78, p < .001, ηp^2^ = .74). Age and Hormone did not interact to influence flank gland diameter *(F* (1, 37) = 1.40, p = .246, ηp^2^ = .04). As such, Hormone significantly increased flank gland diameter in both prepubertal (*Mean Difference* = 0.92 mm, CI 95% [0.26 - 1.56 mm], p < .007, ηp^2^ = .18 and adult subjects *Mean Difference* = 1.45 mm, CI 95% [0.81 - 2.10 mm], p < .001, ηp^2^ = .36).

### 3.2 Flank Marking Behavior

Flank marking data were not normally distributed across groups and required analysis by the Kruskal-Wallis test (Figure 3A). In the scent test, flank marking levels significantly differed between groups (*H* (3, 44) = 11.220, p = .011), such that testosterone-treated adults displayed significantly more flank marking than vehicle-treated adults (p=.01), and both Prepub+0 (p = .003) and Prepub+T (p = .01) groups. In the social interaction test, a similar pattern of group differences was observed (Figure 3B; *H* (3, 42) = 31.36, p < .001). Testosterone-treated adult subjects flank marked more often than Adult+0 (p < .001), Prepub+0 (p < .001), and Prepub+T subjects (p < .001). These data indicate that in either testing context, testosterone activates flank marking behavior in adult but not in prepubertal males.

**Figure 3.**
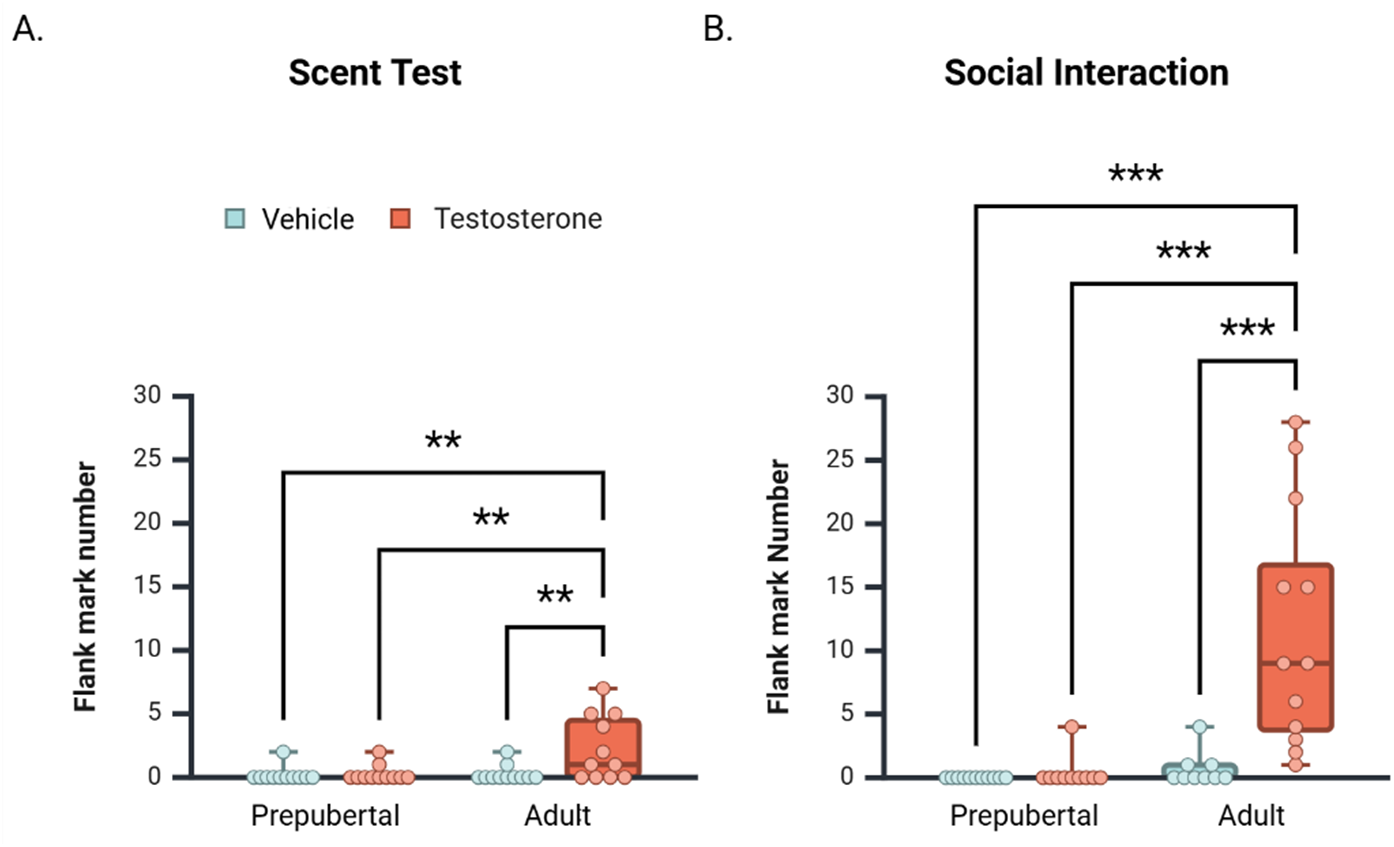
Median number [+/- CI 95%] of flank marks displayed by prepubertal and adult subjects during the Scent Test (A) and Social Interaction Test (B). In both tests, testosterone treatment increased flank marking behavior only in adult subjects. **\*\*** denotes p < .01, *** = p < .001.

### 3.3 Close Contact with Opponent

Prepubertal subjects spent significantly more time in close contact with opponents than did adults (Figure 4A; *F* (1, 38) = 23.70, p < .001, ηp^2^ = .38). An interaction between Age and Hormone also affected total time in close contact (*F* (1, 38) = 6.90, p = .01, ηp^2^ = .15). In prepubertal males, testosterone non-significantly increased close contact duration (*Mean Difference T vs. vehicle* = 73.035 seconds, CI 95% [-19.81 - 165.88 seconds], p = .12). In contrast, testosterone treatment significantly decreased contact duration between adult subjects and their opponents (Figure 4A; *Mean Difference T vs. vehicle* = -97.650 seconds, CI 95% [-191.354 - - 3.946 seconds], p = .04).

**Figure 4.**
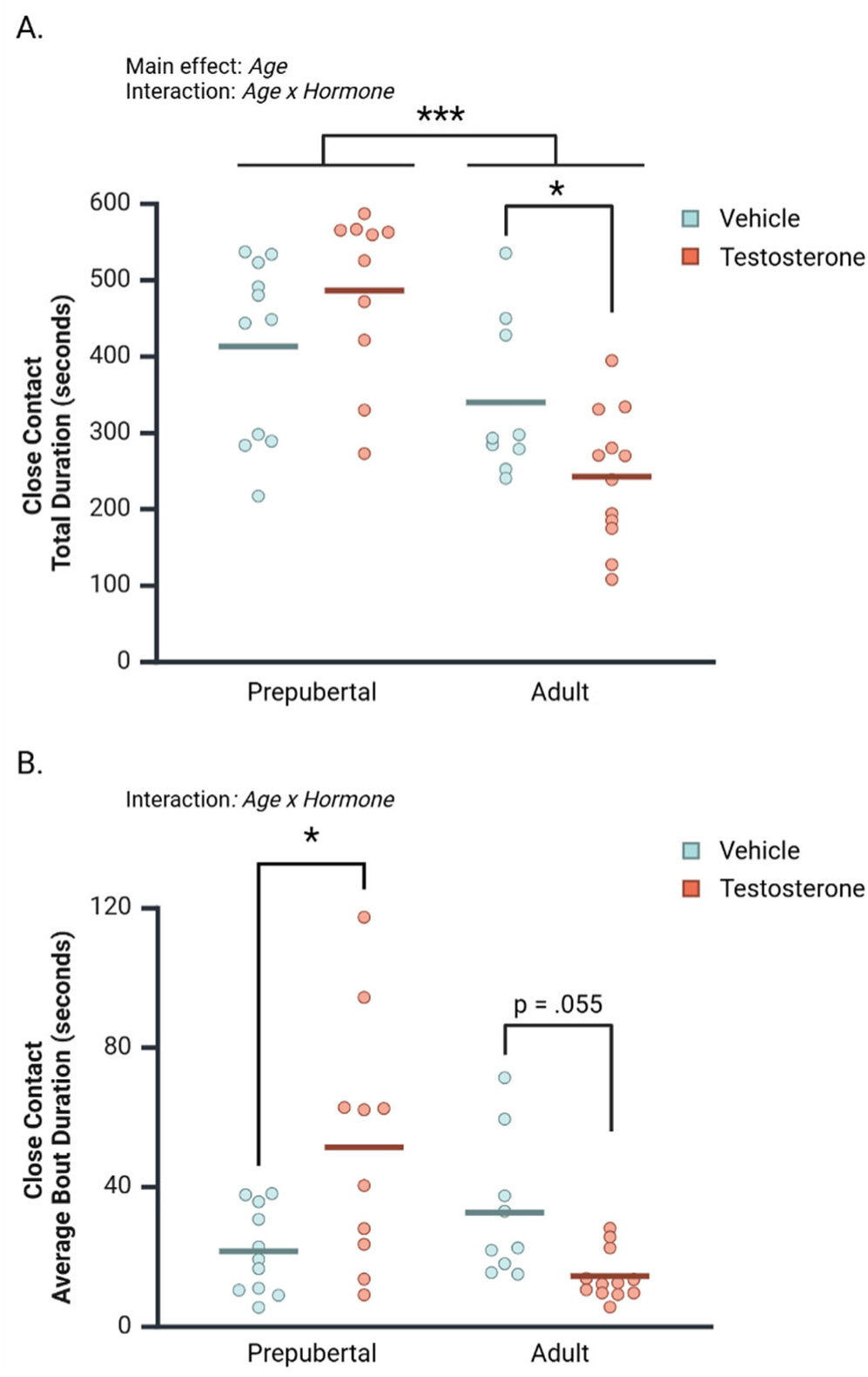
The effects of Age and Hormone on the duration of close contact between subjects and opponents during the Social Interaction Test. **A.** Prepubertal subjects spent significantly more time in close contact with opponents than did adults. A significant interaction between Age and Hormone revealed that T treatment significantly decreased close contact time between adult subjects and opponents, but did not affect prepubertal subject’s time in close contact with opponents. **B.** Age and Hormone interacted to affect close contact average bout durations. T-treatment significantly increased close contact bout durations in prepubertal males and decreased close contact bout durations in adult males. Circles represent individual values, and the bars represent the group mean. **\*** denotes p < .05.

The average bout durations of close contact between subjects and their opponents were also examined (Figure 4B). Age and Hormone significantly interacted to influence the duration of close contact bouts (*F* (1, 38) = 13.78, p < .001, ηp^2^ = .27). This interaction was driven by a significant T-dependent *increase* in contact bout duration in prepubertal subjects (*Mean Difference (T vs. vehicle)* = 29.81 seconds, CI 95% [11.38 - 48.31], p = .002), and a T-dependent *decrease* in contact bout duration in adult subjects (*Mean Difference T vs. vehicle* = -18.26 seconds, CI 95% [-36.893 - .38], p = .055).

### 3.4 Aggressive Displays

Offensive postures significantly differed between treatment groups (Figure 5A; *H* (3, 42) = 11.03, p = .01). Vehicle-treated prepubertal males displayed significantly more offensive postures than Adult+T subjects (p < .001). Offensive postures did not significantly differ between Prepub+0 and Prepub+T groups (p=.06), or between Adult+0 and Adult+T groups (p=.23). Treatment group differences were also observed for paws-on displays (Figure 5B; *H* (3, 42) = 9.42, p = .02). Testosterone-treated adults displayed significantly more paws-on behavior vehicle-treated adults (p = .02) and vehicle-treated prepubertal males (p < .01). Paws-on displays did not significantly differ between Prepub+0 and Prepub+T subjects (p = .08). Attacks on opponents significantly differed between treatment groups (Figure 5C; *H* (3, 42) = 15.60, p = .001). Testosterone-treated prepubertal subjects attacked their opponents more often than vehicle-treated prepubertal subjects (p = .02), and both Adult+0 (p < .001) and Adult+T subjects (p < .01).

**Figure 5.**
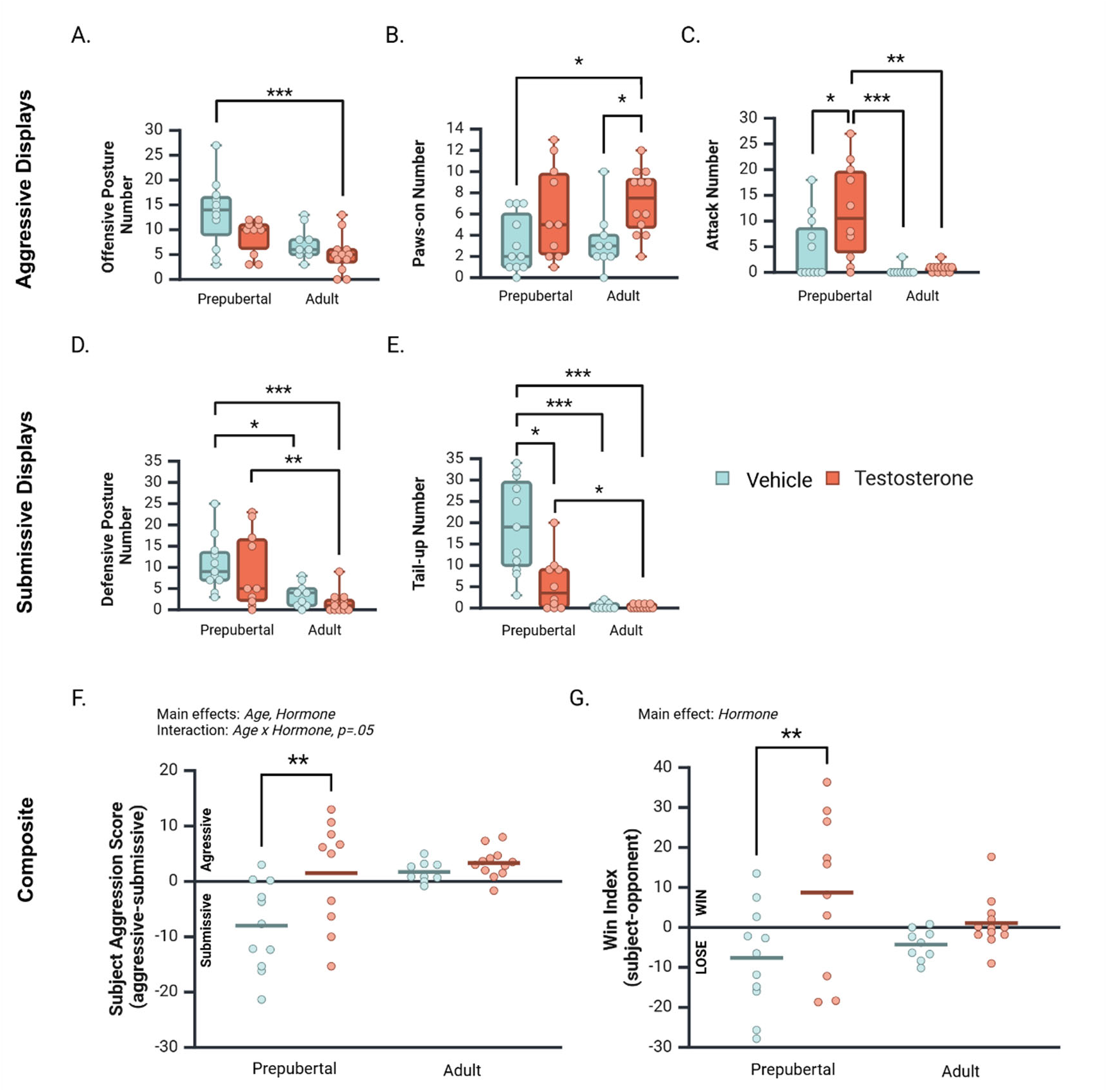
Median number [+/- CI 95%] of aggressive and submissive behaviors displayed by vehicle- and T-treated prepubertal and adult subjects during a social interaction with an age-matched opponent (A-E), and composite mean aggression and win index scores (F-G). **A**. Vehicle-treated prepubertal subjects displayed the highest number of offensive postures, significantly higher than T-treated adults. **B**. Testosterone significantly increased paws-on displays in adult but not prepubertal subjects. **C.** Testosterone significantly increased attacks in prepubertal but not adult subjects. Prepub+T males attacked opponents more often than both Adult+0 and Adult+T groups. **D.** Testosterone did not influence defensive postures of prepubertal or adult subjects, however, prepubertal subjects displayed higher levels of defensive postures than adults. **E**. Tail-up displays were significantly decreased by testosterone in prepubertal but not in adult subjects. Prepubertal subjects displayed higher levels of tail-up displays than adults. **F**. The subject’s aggression score reflects the difference between their average aggressive and submissive displays. Aggression scores were influenced by Age, Hormone, and an interaction between Age and Hormone. Testosterone significantly increased aggression scores in prepubertal males, but not in adult males. **G.** Win index values reflects the difference between aggression scores of subjects and their opponents. Testosterone treatment significantly increased dominance scores overall, and particularly in prepubertal males. * = p < .05, ** = p < .01, *** = p < .001.

### 3.5 Submissive Displays

The distribution of defensive postures significantly differed between treatment groups (Figure 5D; *H* (3, 42) = 16.80, p < .001). Although no difference was observed between vehicle- and T-treated prepubertal groups (p = .25), vehicle-treated prepubertal subjects displayed significantly more defensive postures than both Adult+0 (p < .02) and Adult+T groups (p < .001). In addition, Prepub+T subjects displayed significantly more defensive postures than Adult+T subjects (p < .01; Figure 5D). The distribution of tail-up postures also significantly differed between treatment groups (Figure 5E; *H* (3, 42) = 26.30, p < .001). Prepub+0 subjects displayed significantly more tail-up postures than Prepub+T (p < .02), Adult+0 (p < .001), and Adult+T (p < .001) subjects. In addition, Prepub+T subjects displayed significantly more tail-up postures than did Adult+T subjects (p < .05), but not Adult+0 subjects (p = .07).

### 3.6 Subject Aggression Score and Win Index

Subject aggression scores (average of aggressive displays – average of submissive displays) were significantly higher in adults than in prepubertal subjects (Figure 6A; *F* (1, 38) = 8.40, p < .01, ηp^2^ = .18). In addition, a main effect of Hormone indicated that testosterone treatment increased subject aggression scores overall (*F* (1, 38) = 7.74, p < .01, ηp^2^ = .17). These main effects were qualified by a marginal interaction between Age and Hormone (*F* (1, 38) = 3.90, p < .05, ηp^2^ = .09). Simple comparisons of the effects of testosterone at each age revealed that testosterone treatment significantly increased aggression scores of prepubertal (Figure 6A; *Mean Difference (T vs. vehicle)* = 9.50, CI 95% [3.8 – 15.20], p <.01), but not adult subjects (*Mean Difference (T vs. vehicle)* = 1.62, CI 95% [-4.12 – 7.35], p = .57). Notably, most adult subject’s aggression scores were positive, indicating they displayed more aggressive than submissive behaviors during social interactions, irrespective of testosterone treatment.

A subject win index was calculated to investigate whether testosterone treatment increases the likelihood subjects exert dominance over an opponent before and after adolescence (*subject* aggression score - *opponent* aggression score). Two-factor ANOVA revealed that testosterone treatment significantly increased the win index of subjects compared to vehicle treatment (Figure 6B; *F* (1, 38) = 7.92, p < .01, ηp^2^ = .17). This main effect appeared driven by the difference between Prepub+T and Prepub+0 subjects (*Mean Difference* = 16.34, CI 95% [5.36 – 27.32], p < .01), more so than by the difference between Adult+T and Adult+0 subjects (*Mean Difference* = 5.35, CI 95% [-5.73 - 16.43], p = .34). Age did not significantly impact the win index (Figure 6B; *F* (1, 38) = .31, p = .58, ηp^2^ = .01), nor did Age and Hormone significantly interact (*F* (1, 38) = 2.03, p = .16, ηp^2^ = .05).

## 4.0 Discussion

The current study investigated the effects of testosterone before and after adolescence on agonistic behavioral displays and achieving dominance over an opponent. Our hypothesis that behavioral neural circuits acquire responsiveness to testosterone during adolescence predicted that testosterone would modulate agonistic behavioral displays in adult but not prepubertal males. Flank marking behavior, but no other agonistic behavioral displays, followed the predicted data pattern. During both social interaction and scent tests, testosterone facilitated flank marking behavior *only* in adult subjects, indicating that behavioral responsiveness to testosterone requires developmental processes occurring during adolescence. For all other agonistic behaviors assessed during social interactions, unique behavioral responses to testosterone were observed between prepubertal and adult subjects. Overall, testosterone increased social contact time in prepubertal males and decreased social contact time in adults. Although testosterone increased paws-on investigation in adults, no other aggressive or submissive displays were influenced by testosterone in adults. In prepubertal subjects, testosterone increased attacks and decreased tail-up submissive displays. Testosterone also facilitated prepubertal subject dominance over an opponent (win index). Taken together, these data highlight remarkable adolescent changes in testosterone’s influence on the formation of dominance relationships.

In adult male rodents, the influence of testosterone on aggressive and submissive behaviors is heavily dependent upon the social context (Solomon et al., 2009; Whitten et al., 2024). The current study utilized a neutral arena rather than the subject’s home cage so that defense of a home territory did not bias dominance contests toward subjects. In addition, to permit comparisons of subject agonistic behavior across age, all opponents were testosterone-treated to avoid the potential confounding influence of differing endogenous testosterone levels between prepubertal and adult opponents (Solomon et al., 2009). Previous studies investigating age-related changes in aggressive behavior demonstrate that adolescence is associated with a decrease in attack frequency and a change in the bodily targets of attack (Cervantes et al., 2007; Pellis & Pellis, 1988a, 1988b; Wommack et al., 2003). These previous studies tested gonad-intact male subjects in both neutral and home territories across time. Our findings using a cross-sectional design (before and after adolescence) and testing subjects in a neutral arena support previous reports of an age-related decline in attack frequency. However, to our knowledge, we are the first to report that testosterone *facilitates* attacks in prepubertal males. Although this finding conflicts with a previous report that a 1-wk testosterone treatment fails to influence prepubertal attacks on intruders in their home cage (Romeo et al., 2003), differences in both the duration of testosterone treatment (1 vs. 2 weeks) and the testing environment (home territory vs. neutral arena) likely explain the differential effects of testosterone on attacks between these studies. We also report here that testosterone treatment decreased tail-up submissive postures in prepubertal males. In contrast, adults displayed very few tail-up or defensive postures, irrespective of testosterone treatment. The overall higher levels of submissive behavior displayed by prepubertal males suggests that submissive behavioral displays decrease during adolescent development, much like the well-documented decrease in attacks across the pre- to post-adolescent period (reviewed in Delville et al., 2003).

The neural mechanisms underlying prepubertal and adult differences in behavioral responsiveness to testosterone are currently unknown. Although prepubertal males were more responsive than adults to the effects of testosterone on attacks and tail-up displays, it is unclear whether the lack of responsiveness in adult subjects reflects an adolescent decrease in neural responsiveness to testosterone during adolescent development. Interestingly, gonadectomized and testosterone-treated prepubertal males display higher densities of androgen receptor (AR) than adults within regions of the social behavior network that regulate reproductive and aggressive behavior (Meek et al., 1997). Specifically, AR densities are higher in prepubertal than adult males within the medial amygdala (MeA), medial preoptic nucleus (MPNmag), and bed nucleus of the stria terminalis (BNST). Thus, it is possible that adolescent decreases in androgen receptor within this network contributes to the differences in behavioral responsiveness to testosterone observed between prepubertal and adult males. Given that the effects of testosterone on aggression are mediated by social context in adulthood (Oliveira, 2009; Solomon et al., 2009; Whitten et al., 2024), it is also likely that differences observed between prepubertal and adult males relate to adolescent development of corticolimbic circuits underlying social cognition (Blakemore, 2008; Murray et al., 2024) and threat perception (Spielberg et al., 2015). Adolescent changes in corticolimbic circuits promote the context-appropriate expression of social behavior (K. De Lorme et al., 2013; K. C. De Lorme & Sisk, 2013), and may also change the context-dependent effects of testosterone on aggressive and submissive behavioral displays in adulthood.

Flank marking is a testosterone-dependent form of scent communication utilized by adult male Syrian hamsters to communicate dominance status (Ferris et al., 1987; Johnston, 1981). Since the effects of testosterone on behavior are often context-dependent, we evaluated flank marking behavior in two different contexts: during social interactions and in response to male odors alone. In both contexts, testosterone treatment stimulated flank marking behavior in adult males, but not in prepubertal males. This suggests that testosterone’s effects on flank marking behavior do not appear until after puberty and adolescence. We also evaluated whether peripheral flank glands were responsive to the presence of testosterone prior to puberty and found that testosterone treatment increased flank gland diameters in both prepubertal and adult males. Therefore, the lack of behavioral response to testosterone in prepubertal males is not likely due to insensitivity of peripheral flank gland tissues. Instead, the inability of testosterone to activate flank marking behavior before adolescence is likely due to unresponsive behavioral neural circuits at this age. Indeed, exposure to gonadal hormones, specifically during adolescence, is necessary for the activation of flank marking behavior by testosterone in adulthood (Schulz et al., 2006). Testosterone’s unique effects on flank marking and aggression may indicate these behaviors have distinctive underlying neural circuits within the social behavior network that are differentially regulated by testosterone before and after puberty.

To investigate whether testosterone treatment facilitated dominance over an opponent in prepubertal and adult subjects, we calculated a win index by subtracting the opponent’s aggression score (mean aggressive displays – mean submissive displays) from each subject’s aggression score. Testosterone treatment significantly increased the subject win index overall, and especially in prepubertal males. A win index value of zero indicates that no clear dominance relationship was formed during the social interaction. In prepubertal males, win index values were clearly either above or below zero, indicating a definitive win or loss during the social interaction. In adults, win index values clustered more closely around zero, and in many cases no clear winner or loser emerged from the interaction. Given that aggressive behavior is sensitive to the test environment and tends to be expressed at lower levels in a neutral environment than in a home cage territory (Mink & Adams, 1981), the low level of attacks displayed by adult subjects is likely due to the neutral test arena. Thus, our findings suggest that in a neutral testing environment, testosterone increases competitive motivation in prepubertal but not adult males. These data further support likelihood that the context-appropriate expression of aggressive behavior requires developmental processes occurring during adolescence (K. C. De Lorme & Sisk, 2013).

## 5.0 Conclusions

We provide here evidence that some, but not all aspects of agonistic behavior are sensitive to the activational effects of testosterone prior to adolescence, and that activational effects of testosterone differ substantially between prepubertal and adult males. Several studies in humans have linked early pubertal timing with increased adolescent aggression and externalizing symptoms (Chen & Raine, 2018; Cota-Robles et al., 2002; Ge et al., 2006; Glowacz & Bourguignon, 2015; Najman et al., 2009). Our findings support the possibility that early pubertal gonadal steroid hormones act on responsive developing neural networks and increase risk for externalizing symptoms and aggressive behavior during adolescence.

## 6.0 CRediT Authorship Contribution Statement

**Arthur Castaneda**: Data curation, formal analysis, original draft, visualization, and review and editing. **Conner Whitten**: Data curation, formal analysis, visualization, validation, and review and editing. **Tami Menard**: Methodology, investigation, project administration, and formal analysis. **Cheryl Sisk:** Funding acquisition and review and editing. **Matthew Cooper:** Validation, visualization, and review and editing. **Kalynn Schulz**: Conceptualization, methodology, investigation, project administration, data curation, formal analysis, visualization, review and editing, supervision, and funding acquisition.

